# Autophagy gene-dependent intracellular immunity triggered by interferon-γ

**DOI:** 10.1101/2021.05.10.443539

**Authors:** Michael R. McAllaster, Jaya Bhushan, Dale R. Balce, Anthony Orvedahl, Arnold Park, Seungmin Hwang, Meagan E. Sullender, L. David Sibley, Herbert W. Virgin

## Abstract

Genes required for the lysosomal degradation pathway of autophagy play key roles in topologically distinct cellular processes with significant physiologic importance. One of the first-described of these *ATG* gene-dependent processes is the requirement for a subset of *ATG* genes in interferon-γ (IFNγ)-induced inhibition of norovirus and *Toxoplasma gondii* replication. Herein we identified new genes that are required for or that negatively regulate this immune mechanism. Enzymes involved in the conjugation of UFM1 to target proteins including UFC1 and UBA5, negatively regulated IFNγ-induced inhibition of norovirus replication via effects of *Ern1*. IFNγ-induced inhibition of norovirus replication required *Wipi2b* and *Atg9a*, but not *Becn1* (encoding Beclin1), *Atg14*, or *Sqstm1*. The phosphatidylinositol-3-phosphate and ATG16L1 binding domains of WIPI2B were required for IFNγ-induced inhibition of norovirus replication. Both *WIPI2* and *SQSTM1* were required for IFNγ-induced inhibition of *Toxoplasma gondii* replication in HeLa cells. These studies further delineate the mechanisms of a programmable form of cytokine-induced intracellular immunity that relies on an expanding cassette of essential *ATG* genes to restrict the growth of phylogenetically diverse pathogens.

**Importance:** Interferon-γ is a critical mediator of cell-intrinsic immunity to intracellular pathogens. Understanding the complex cellular mechanisms supporting robust interferon-γ-induced host defenses could aid in developing new therapeutics to treat infections. Here, we examined the impact of autophagy in the interferon-γ induced host response. We demonstrate that CRISPR-Cas9 screens specifically targeting the autophagy pathway uncover a role for WIPI2 in IFNγ-induced inhibition of *Norovirus* replication in mouse cells and IFNγ mediated restriction of the *Toxoplasma gondii* parasitophorous vacuole in human cells. Furthermore, we found perturbation of UFMylation pathway components led to more robust IFNγ-induced inhibition of *Norovirus* due to ER stress *in vitro*. Enhancing or inhibiting these dynamic cellular components could serve as a strategy to weaken intracellular pathogens and maintain an effective immune response.

## Introduction

Macroautophagy (autophagy herein) requires formation of an isolation membrane that envelops cytoplasmic materials, organelles or invading pathogens in a closed double membrane-bound autophagosome. Autophagosomes fuse with lysosomes to facilitate degradation of captured material (1, 2). Autophagy requires a series of autophagy genes (*ATG* genes), many of which are conserved broadly in evolution. It is now clear that these essential *ATG* genes are also required for additional topologically distinct cellular processes that have significant physiologic importance (3, 4, 5). Many of these autophagy-independent, but *ATG* gene-dependent, processes occur in myeloid cells. These include essential roles for *ATG* genes in the anti-microbial action of the key cytokine IFNγ (also referred to as Type II IFN), the deposition of the ATG8-family proteins on the cytoplasmic surface of phagosomes (2, 6), the fusion of lysosomes to the polarized ruffled membrane of osteoclasts to secrete lysosomal proteases for extracellular degradation of bone (7), the regulation of neutrophilic inflammation during *Mycobacterium tuberculosis* infection (8) and the inhibition of inflammatory activation of tissue-resident macrophages (9). Of particular note, the role of *ATG* genes in the actions of IFNγ are fundamentally important to survival of mammals because this cytokine plays an essential role in triggering cell-intrinsic immunity to intracellular bacteria, viruses and parasites.

The fact that *ATG* genes play roles in both autophagy and topologically distinct cellular processes makes it imperative to further define the molecular machinery that is in common to, or distinguishes, these distinct cellular events. Here we focus on IFNγ-induced *ATG* gene-dependent intracellular immunity to murine norovirus (norovirus herein) and *Toxoplasma gondii* (*T. gondii)*. Inhibition of intracellular replication of these pathogens requires *ATG* genes *Atg7, Atg5, Atg16l1, Atg12, Atg3* and ATG8 family members (10–18). ATG3, ATG7 and the ATG5-ATG12-ATG16L1 (herein ATG5-12-16L1) complex are required for the ubiquitin-like conjugation system of autophagy that conjugates phosphatidyl-ethanolamine (PE) to localize ATG8 proteins on membranes (1, 4). However, the topology of the intracellular events triggered by IFNγ are distinct for the two pathogens. *T. gondii* replicates in a parasitophorous vacuole sequestered away from the cytoplasm, while norovirus replicates on the cytoplasmic face of membranes. In mouse, IFNγ-induced *ATG* gene-dependent clearance of *T. gondii* occurs by destruction of the parasitophorous vacuole (10, 13), while in human cells this pathway limits replication but does not eliminate the parasite (15, 19). In contrast, IFNγ inhibits norovirus replication by disrupting the formation and/or function of the membranous replication compartment upon which viral RNA synthesis occurs (10–18).

Here, we show that an immortalized murine macrophage-like microglial cell line (BV-2 cells) (20) recapitulates *ATG* gene-dependent events in IFNγ-induced inhibition of norovirus replication as originally described in primary macrophages. Using CRISPR-Cas9 screening, we identified additional positive and negative regulators of this process. *Ufc1* and *Uba5* regulated IFNγ inhibition of norovirus replication. In contrast, *Atg9a* and *Wipi2b* were required for efficient IFNγ-induced inhibition of norovirus replication. The ATG16L1-and phosphatidylinositol 3-phosphate (PtdIns(3)P)-binding domains of WIPI2B were required for efficient IFNγ-induced inhibition of norovirus replication. *WIPI2* was also required for IFNγ-induced inhibition of *T. gondii* replication in human cells. In contrast to findings in the norovirus system, *SQSTM1* (herein *P62)* was required for IFNγ-induced inhibition of *T. gondii* replication. Thus, components of the autophagy machinery outside of the ubiquitin-like conjugation systems involving the ATG5-12-16L1 complex are important components of a cellular immune mechanism used by IFNγ to block intracellular pathogen replication, potentially identifying targets for modulation of this type of immunity.

## RESULTS

### IFNγ inhibits norovirus replication in BV-2 cells in a *Stat1*- and *Irf1*-dependent manner

BV-2 cells are permissive for replication of murine norovirus strain MNoV^CW3^ (21), allowing us to test whether, in these cells, norovirus replication was inhibited by recombinant IFNγ as measured by plaque assay and cellular ATP content as a proxy for cell viability (Fig. S1A). Pretreatment of BV-2 cells with IFNγ decreased MNoV^CW3^ replication ∼5000-fold (Fig. 1A) and cytopathicity by 50% (Fig. 1B). *Stat1* and *Irf1* encode transcription factors essential for IFNγ responses (22), and are required for robust IFNγ-induced inhibition of norovirus replication both *in vitro* in primary macrophages and *in vivo* (23–25). To determine whether these transcription factors were required for IFNγ-induced inhibition of MNoV^CW3^ replication in BV-2 cells we generated two independent clonal *Stat1* or *Irf1* knockout BV-2 cell lines (*Stat1*^*-/-*^, *Irf1*^*-/-*^)(Fig. 1A and B). Throughout this work we confirmed deletion of genes in cell lines using next generation sequencing (Table S1). IFNγ failed to efficiently inhibit norovirus replication and cytopathicity in either *Stat1*^*-/-*^or *Irf1*^*-/-*^cells (Fig. 1A, 1B), faithfully replicating the requirement for these genes in primary cells.

**Figure 1:**
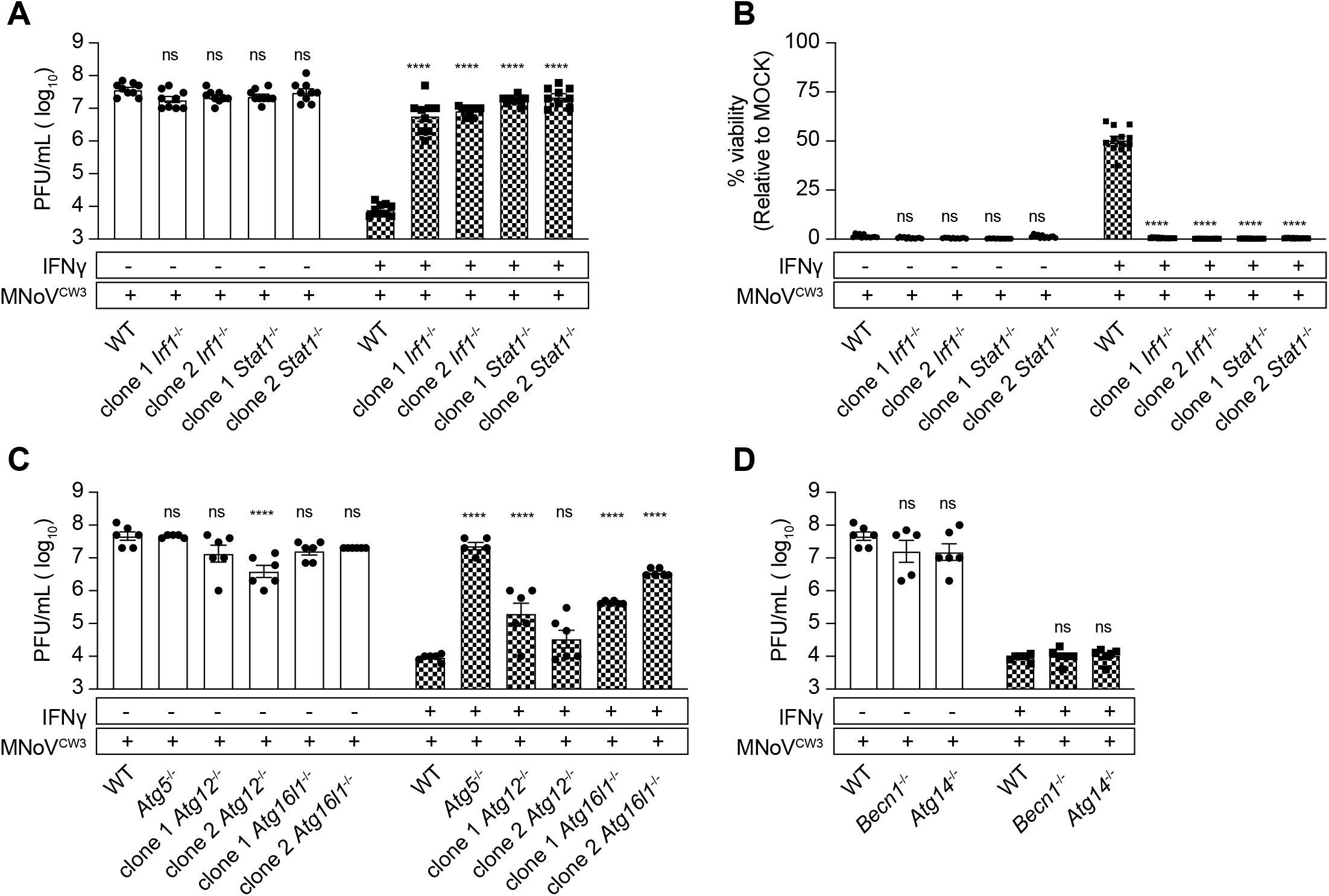
*Stat1, Irf1, Atg5, Atg12* and *Atg16l1* are required for IFNγ-induced inhibi-tion of norovirus replication in BV-2 cells, *Atg14* and *Becn1* are not. (A) Plaque assay of WT, *Irf1*^-/-^and *Stat1*^-/-^BV-2 cells as described in (Fig. S1). (B) Viability assay of WT, *Irf1*^-/-^and *Stat1*^-/-^BV-2 cells as described in (Fig. S1). (C) Plaque assay of WT, *Atg5*^-/-^, *Atg12*^-/-^and *Ag16l1*^-/-^BV-2 cells as described in (Fig. S1). (D) As in (Fig. S1) for WT, *Atg14*^-/-^and *Becn1*^-/-^BV-2 cells. Average data pooled from 2-3 independent experi-ments are represented as means ± SEM. *P* value ≤ 0.05 (*), ≤ 0.01(**), ≤0.001 (***), ≤ 0.0001 (****) were considered statistically significant. ns, not significant. *P* value deter-mined by 2-way ANOVA with Dunnett’s multiple comparison test.

### The ATG5-12-16L1 complex is required for IFNγ-induced inhibition of norovirus replication in BV-2 cells

The ATG5-12-16L1 complex is crucial for IFNγ-induced control of MNoV^CW3^ in bone marrow-derived murine macrophages (11). Using *Atg5* knockout BV-2 cells (*Atg5*^-/-^) (26, 27) we confirmed that *Atg5* was required for IFNγ-induced inhibition of MNoV^CW3^ replication (Fig. 1C). To further define the role of the ATG5-12-16L1 complex we generated two independent clonal knockout BV-2 cell lines for each of *Atg12* and *Atg16l1* (*Atg12*^*-/-*^, *Atg16l1*^*-/-*^) (Table S1)(27). IFNγ failed to efficiently inhibit MNoV^CW3^ replication in cells lacking either *Atg12* or *Atg16l1* (Fig. 1C). While there were significant decreases IFNγ inhibition of norovirus replication in cells lacking *Atg12* or *Atg16l1*, there was variation between clonal knockout cell lines. Deep sequencing confirmed saturating indel frequencies in both *Atg12* and *Atg16l1* in clonal knockout cells (Table S1) supporting a role for *Atg12* and *Atg16l1* in IFNγ inhibition of MNoV^CW3^ replication in these cells.

### The *ATG* genes *Becn1* and *Atg14* are not required for IFNγ-induced inhibition of norovirus replication

Certain upstream components of the autophagy pathway are not required for IFNγ-induced inhibition of norovirus or *T. gondii* replication (10, 11, 15, 16). To confirm these observations in BV-2 cells, we determined whether IFNγ efficiently inhibits MNoV^CW3^ replication in clonal *Atg14*^-/-^BV-2 cells (26, 27) and clonal *Becn1*^-/-^cell lines (27). Neither *Atg14* nor *Becn1* were required for IFNγ-induced inhibition of MNoV^CW3^ replication (Fig. 1D). Together with the data on the role of *Stat1, Irf1, Atg5, Atg12 and Atg16l1* above these data support the validity of BV-2 cells as a model to further define mechanisms of IFNγ-induced *ATG* gene-dependent immunity to norovirus.

### CRISPR screen design for identification of genes involved in IFNγ-induced inhibition of norovirus cytopathicity in BV-2 cells

We used CRISPR screening to determine the possible contribution of genes selected for their potential roles in autophagy or other *ATG* gene-dependent cellular processes to IFNγ action. We designed an autophagy sgRNA library containing 1 to 4 independent guides targeting 695 candidate genes (Table S2). We included 300 guides with no known target as controls (2979 guides total) (28). Genes were selected using three criteria: (i) present in the autophagy interaction landscape, including the baits used, as defined by Behrends et al. (29); (ii) murine genes related to autophagy using GO ontology (30, 31); (iii) murine genes corresponding to human genes related to autophagy using GO ontology (30, 31).

To enhance the sensitivity of this CRISPR screen we targeted generation of a cell pool with each guide represented by ∼2000 cells (32). After experimental selection (Fig. S2A), we quantified guides by DNA sequencing and determined the significance of differences in guide frequencies in different comparisons using STARS and negative binomial analysis (21, 26–28, 33) with a threshold of FDR < 0.1 to identify genes for further consideration (27).

### Identification of genes required for cell survival in the absence of norovirus infection

The effects of genes required for cell survival or proliferation may obscure identification of those required for a phenotype, especially when cell death is part of the biology being studied as is the case for norovirus replication and IFNγ treatment (21, 27, 28). We therefore identified differences in guide frequency between the original cell library and cells passaged under mock conditions (Fig. 2A, Table S3) as well as between cells passaged under mock conditions compared to those treated with IFNγ at two doses (Fig. 2B and 2C, Table S5). Mock passage did not enrich guides for any gene but decreased guides for 52 genes, suggesting that these are essential genes for survival of BV-2 cells under these conditions. Among the targeted genes were *Cdc37, Adsl*, and *Cct2*. Mutations in ADSL result in the rare autosomal recessive disorder Adenylosuccinate lyase deficiency (34) while CCT2 mutations evoke the rare disease Leber Congenital Amaurosis (35). We compared these candidate essential genes to those identified in Project Achilles (36–38). Three of 52 genes lacked a human homolog and one did not have specific guides in Project Achilles. Therefore 48 candidate essential genes were considered for this comparison (Table S4). 43 of 48 (95%) depleted genes were considered common essential among immortalized cells (Table S4) (36–38) thus validating our screening approach. We deprioritized further analysis of genes whose guides were depleted under control conditions.

**Figure 2:**
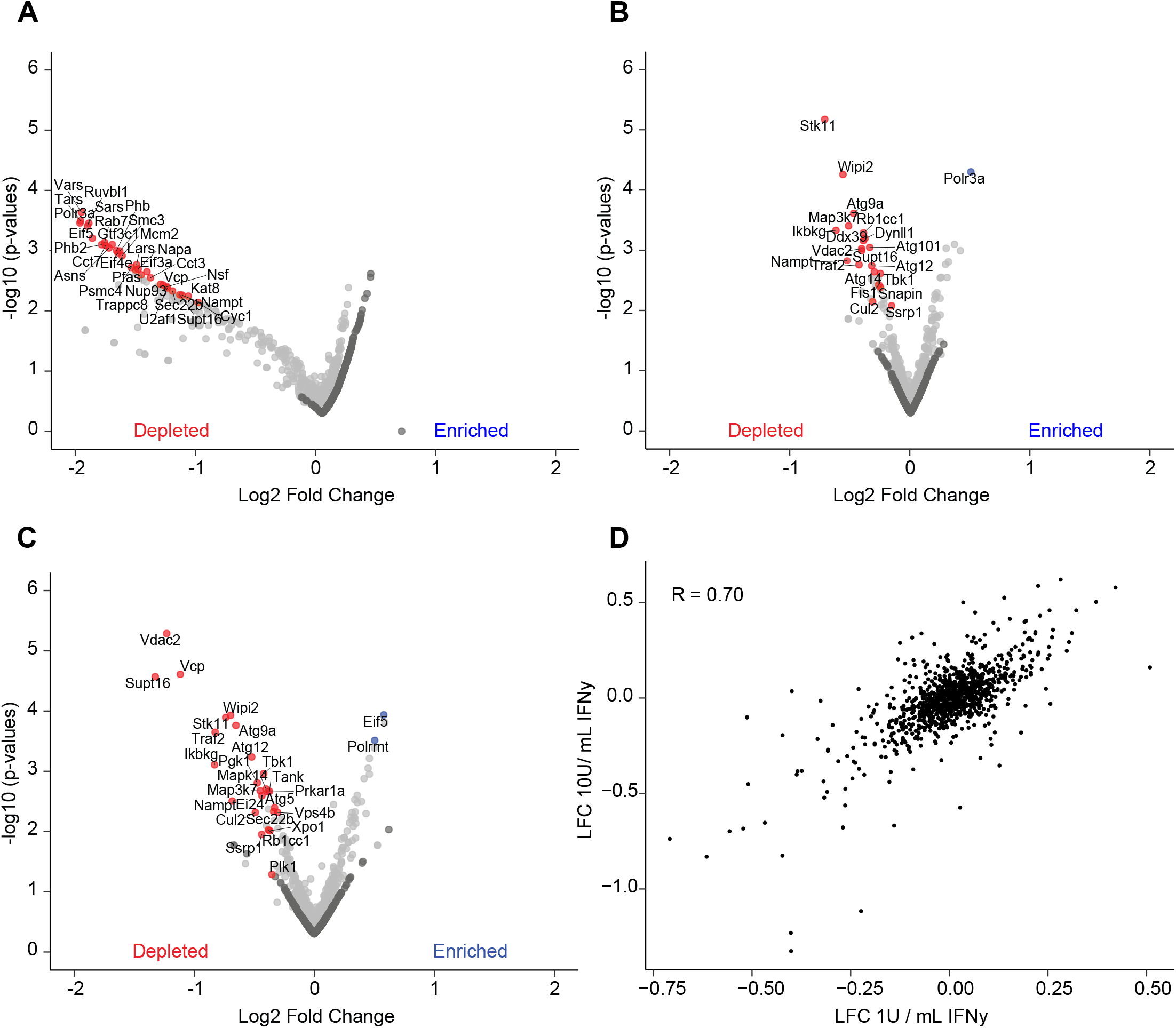
Identification of genes required for viability in passage or IFNγ-treated BV-2 cells. (A) Volcano plot of mock treatment guides enriched or depleted relative to five days post puromycin selection. (B) Volcano plot of guides enriched or depleted after 1U/mL IFNγ treatment relative to mock treatment. (C) as in (B) for 10U/mL IFNγ treatment. (D) Average log2 fold change (LFC) of 1U/mL IFNγ condition versus LFC of 10U/mL IFNγ condition. Pearson correlations are indicated. For volcano plots the LFC of all sgRNAs for each gene is plotted against the –log10(p-value) for each gene. Blue and red highlighted genes in (A) (B) (C) represent a STARS score with FDR < 0.01, dark grey genes represent non-targeting guides.

### Identification of genes enriched or depleted in the presence IFNγ

IFNγ treatment at a dose of 1 U/mL enriched guides for 7 genes (Fig. 2B, Table S5). Escalation of the IFNγ dose to 10 U/mL enriched guides for 20 genes (Fig. 2C, Table S5). Among the targeted genes were *Eif5, Polrmt, Eif3a, Polr3a, Eif4g2* and *Eif4e* (Fig. 2B and 2C, Table S5). These genes regulate mRNA translation and immune cell activation (39–41). IFNγ treatment at 1 U/mL depleted guides for 25 genes (Fig. 2B, Table S5), and at a dose of 10 U/mL depleted guides for 24 genes (Fig. 2C, Table S5). We observed agreement at the gene-level between treatment with 1 U/mL or 10 U/mL IFNγ, with a Pearson’s correlation of 0.70 (Fig. 2D). Among the targeted genes were *Traf2, Tbk1, Ikbkg, Tank, Ei24, Map3k7, Atg5 and Atg14* (Fig. 2B and C, Table S5). *Traf2, Tbk1, Ikbkg, Tank, Ei24 and Map3k7* regulate the tumor necrosis factor receptor signaling pathway (42–44). Consistent with these results, IFNγ-induced cell death in BV-2 cells is mediated by tumor necrosis factor (27). Our results confirmed the critical role for *Atg14* and *Atg5* in inhibiting IFNγ-induced cell death in BV-2 cells (Fig. 2C, Table S5) (27). Guides for *Wipi2, Rb1cc1* (also known as *Fip200*), *Atg9a, Atg101 and Atg12* were also depleted, though these genes were not reported to protect BV-2 cells from IFNγ-induced cell death (Fig. 2B and 2C, Table S5) (27).

### Identification of genes that enhance IFNγ-induced inhibition of norovirus induced cytopathicity

Guides for 18 genes (Fig. 3A, Table S6) were enriched in IFNγ-treated and norovirus infected cells compared to mock conditions. Among the targeted genes were *G3bp1, Sptcl1* and *Sptcl2* which are required for efficient norovirus replication (21, 45–47). Among the genes identified as candidates for enhancing IFNγ-induced inhibition of norovirus replication were *Iqgap1* and *Ufc1* (Fig. 3A, Table S6). Deficiency of IQGAP1 in human monocytic cells results in hyperactive type I IFN responses to cytosolic nucleotides (48). *Ufc1* and *Uba5* encode enzymes in the ubiquitin-like system that covalently conjugates UFM1 to target proteins (UFMylation herein) (26, 49–51). UFMylation is required to maintain ER homeostasis, so that in the absence of UFMylation consequent ER stress can lead to overactivation of IFNγ responses (26). We therefore quantified the effects of IFNγ on norovirus replication in cells lacking *Ufc1* or *Uba5* (*Ufc1*^*-/-*^, *Uba5*^*-/-*^) (26). IFNγ more potently inhibited norovirus replication in these cells (Fig. 3D and E) as shown by diminished potency of IFNγ upon expression of wild type UFC1 (*Ufc1*^WT^ or *Uba5*^WT^), but not the enzymatically dead versions of these proteins (*Ufc1*^ΔC116A^; *Uba5*^ΔC248A^) (Fig. 3D and E) (26). UBA5 contains a UFM1 interaction motif disrupted upon the mutations of W340A/L344A (*Uba5*^ΔUFIM^) (26, 49, 50). Disruption of this motif prevented UBA5 from rescuing the increased IFNγ potency observed in *Uba5*^*-/-*^cells (Fig. 3E). Consistent with data from Balce *et al*., *Ern1* deletion reversed effects on IFNγ potency observed in *Uba5*^*-/-*^cells, consistent with a role for this aspect of the ER stress response in modulating IFNγ action (Fig. 3F) (26).

**Figure 3:**
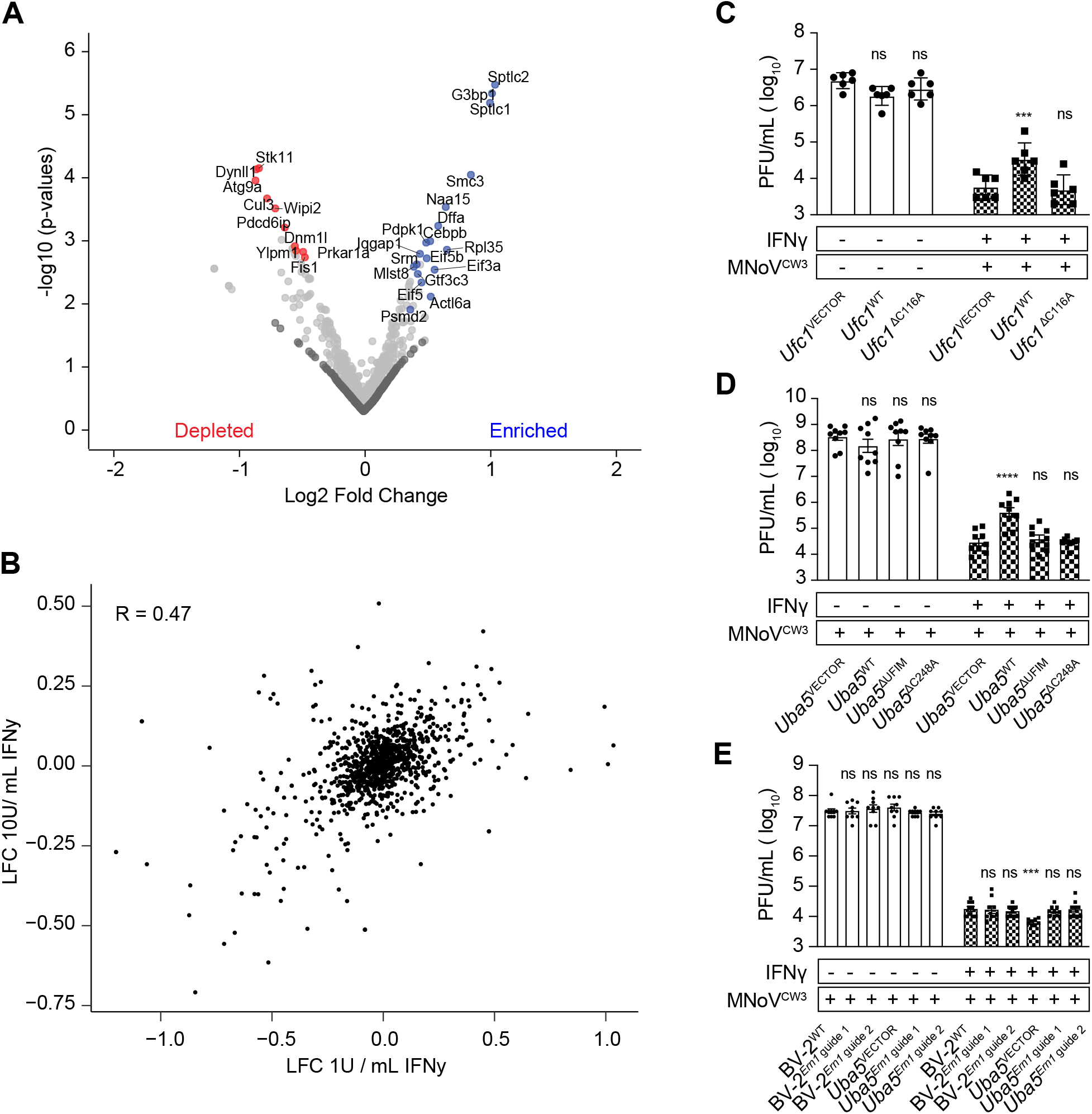
Identification of genes required for of IFNγ-induced norovirus cytopathicity in BV-2 cells. (A) Volcano plot of guides enriched or depleted after 1U/mL IFNγ + MNoV^CW3^ infection relative to mock treatment. (B) Average LFC of 1U/mL IFNγ condition versus LFC of 1U/mL IFNγ + MNoV^CW3^ condition. (C) Plaque assay of *Ufc1*^VECTOR^, *Ufc1*^WT^, *Ufc1*^ΔC116A^ BV-2 cells as described in (Fig. S1). (D) Plaque assay of *Uba5*^VECTOR^, *Uba5*^WT^, *Uba5*^ΔUFIM^, *Uba5*^ΔC248A^ BV-2 cells as described in (Fig. S1). (E) Plaque assay of BV-2^WT^, BV-2^Ern1 guide 1^, BV-2^Ern1 guide 2^, *Uba5*^VECTOR^, *Uba5*^Ern1 guide 1^, *Uba5*^Ern1 guide 2^ as described in (Fig. S1). For volcano plots the aver-age log2 fold change (LFC) of all sgRNAs for each gene is plotted against the –log10(p-value) for each gene. Blue and red highlighted genes in (A) represent a STARS score with FDR < 0.01, dark grey genes represent non-targeting guides. Values in (C) (D) (E) represent means ± SEM from two to three independent experiments. *P* value ≤ ≤0.001 (***), 0.0001 (****) were considered statistically significant. ns, not significant. *P* value determined by 2-way ANOVA with Dunnett’s multiple comparison test.

### Identification of candidate genes required for efficient inhibition of norovirus-induced cytopathicity by IFNγ

We reasoned that guides depleted after pre-treatment with 1U/mL IFNγ and infection with norovirus compared to mock conditions might represent genes required for IFNγ-induced inhibition of norovirus infection. We compared guide frequencies between mock and 1U/mL IFNγ + MNoV^CW3^ infection, reasoning that guides decreased in cells surviving MNoV^CW3^ infection of IFNγ-treated cells might have a role in IFNγ-induced inhibition of MNoV^CW3^ replication (Fig. 3A). We observed intermediate agreement of gene-level results, with a Pearson’s correlation of 0.45 between 1 U/mL IFNγ treatment alone compared to 1U/mL IFNγ + MNoV^CW3^ infected cells (Fig. 3B). Guides for 32 genes (Fig. 3A, Table S6) were decreased in frequency in IFNγ + MNoV^CW3^ infected cells. *Wipi2* and *Atg9a* had not been previously identified as required for *ATG* gene dependent innate immunity. We generated one clonal BV-2 cell line lacking *Wipi2* (*Wipi2*^-/-^; clone 1) (Table S1) and one heterozygous cell line lacking *Wipi2* (*Wipi2*^-/+^ ; clone 2) (Table S1). We also examined BV-2 cells lacking *Atg9a* (*Atg9a*^-/-^) (Table S1) (26). *Wipi2* and *Atg9a* were required for efficient IFNγ-induced inhibition of MNoV^CW3^ replication (Fig. 4A), indicating that they constitute key components of the intracellular mechanisms used by IFNγ to inhibit norovirus infection.

**Figure 4:**
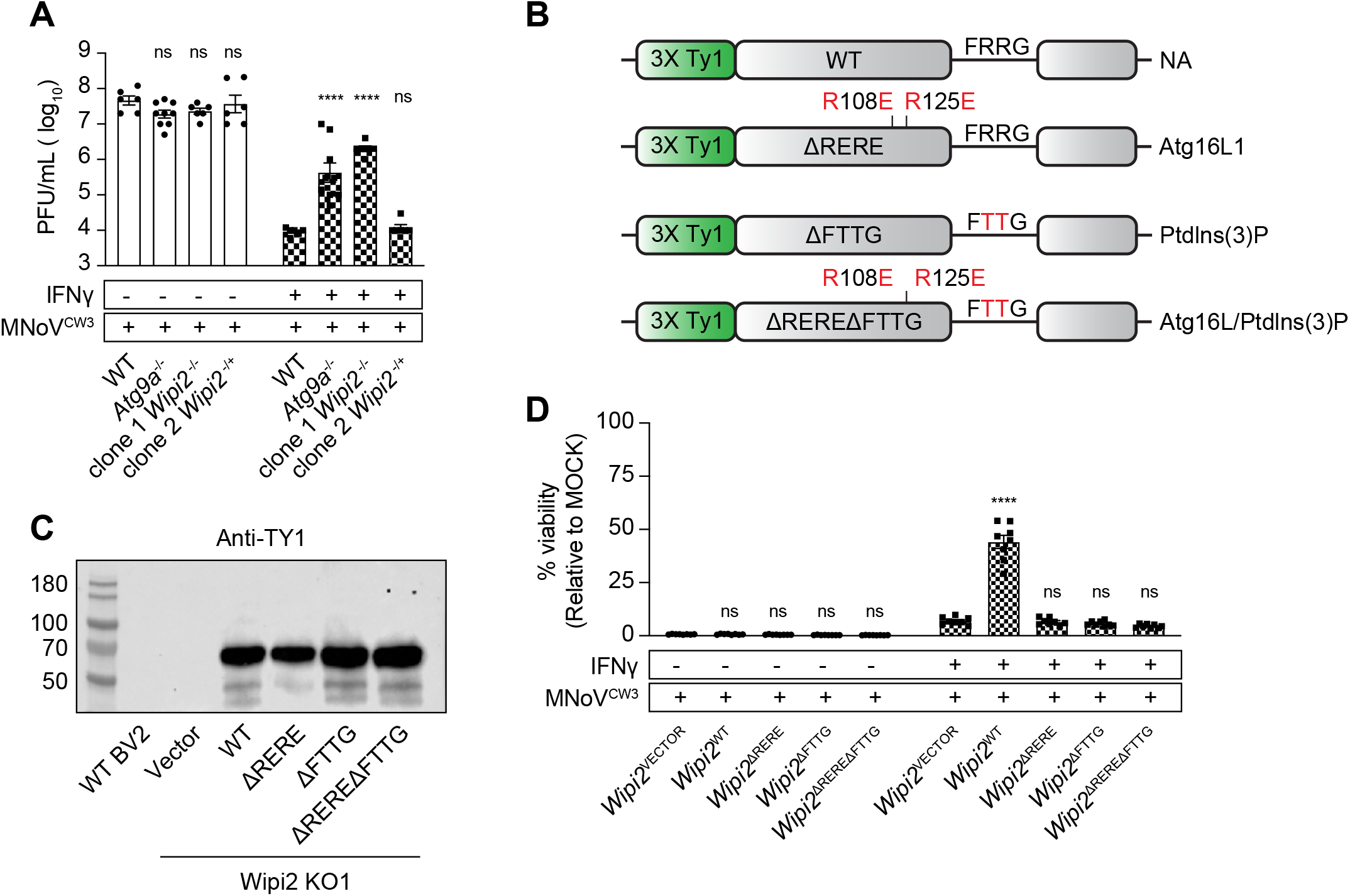
ATG16L1 and PtdIns(3)P binding domains of WIPI2B are required for IFNγ-induced inhibition of norovirus cytopathicity in BV-2 cells. (A) Plaque assay of WT, *Atg9a*^-/-^and *Wipi*2^-/-^BV-2 cells. (B) Schematic of WIPI2B complementation in clone 1 *Wipi2*^-/-^BV-2 cells. (C) Western blot detection of TY1 epitope in WIPI2B complemented cells. (D) Viability assay of *Wipi2*^VECTOR^, *Wipi2*^WT^, *Wipi2*^ΔRERE^, *Wipi2*^ΔRERE^ and *Wipi2*^ΔREREΔFTTG^ BV-2 cells as described in (Fig. S1). Values in (A) and (D) represent means ± SEM from two to three independent experiments. *P* value ≤ 0.05 (*), ≤ 0.01(**), ≤0.001 (***), ≤ 0.0001 (****) were considered statistically significant. ns, not significant. *P* value determined by 2-way ANOVA with Dunnett’s multiple comparison test.

### ATG16L1 and PtdIns(3)P binding domains of WIPI2 are required for IFNγ-induced inhibition of norovirus replication

We analyzed the role of WIPI2 in greater detail because it interacts with ATG16L1 (52, 53), which plays a key role in IFNγ-induced inhibition of norovirus replication (Fig. 4B) (11, 16). We defined the effects of the R108E/R125E (ΔRERE) mutations that inhibit binding to ATG16L1, the R224T/R225T (ΔFTTG) mutations that inhibit binding to PtdIns(3)P, and the double mutant ΔRERE/ΔFTTG (Fig. 4B) (52). We stably expressed these proteins and wild type WIPI2B in *Wipi2*^-/-^(clone 1) cells and assessed protein expression by western blot analysis (Fig. 4C), probing for an N-terminal epitope tag, revealing equivalent expression of the different constructs (Fig. 4C). Expression of WIPI2B rescued the capacity of IFNγ to inhibit norovirus infection in *Wipi2*^-/-^cells (Fig. 4D). In contrast, neither WIPI2B^ΔRERE^ nor WIPI2B^ΔFTTG^ rescued inhibition of MNoV^CW3^ after IFNγ-treatment (Fig. 4D). The double mutant WIPI2B^ΔREREΔFTTG^ protein also failed to restore IFNγ activity (Fig. 4D). Thus WIPI2B binding to both ATG16L1 and PtdIns(3)P was necessary for immune control of norovirus by IFNγ.

### Determining the generality of a role for *WIPI2* in IFNγ-induced *ATG* gene-dependent immunity

The original reports of IFNγ-induced *ATG* gene-dependent immunity demonstrated targeting of both norovirus, a cytoplasmic RNA virus, and the apicomplexan parasite *T. gondii* (10, 11). To determine whether *WIPI2* plays a more general role in IFNγ-induced immunity, we quantitated *T. gondii* replication within the parasitophorous vacuole (PV) in parental and two independent *WIPI2*^*-/-*^human HeLa cells (Table S1), using a type III parasite that is susceptible to *ATG* gene-dependent IFNγ-induced growth control (15, 19). *T. gondii* multiplies by binary fission with a half-life of ∼8 hours, generating vacuoles of different sizes containing clusters of 1 to ≥ 8 parasites over 24 hours. Parasitophorous vacuoles become labeled by ubiquitin in IFNγ-treated cells (15). Ubiquitin-positive vacuoles are targeted in an *ATG* gene-dependent manner resulting in smaller vacuoles and decreased numbers of parasites per vacuole (15), we therefore counted parasites per ubiquitin-positive vacuole in wild type and *WIPI2*^*-/-*^HeLa cells (Fig. 5A) (15). In wild type cells, we did not observe a growth restriction phenotype in PVs lacking ubiquitin (Fig. 5A, left), however most ubiquitin-positive vacuoles contained only 1 parasite, indicative of IFNγ-induced growth restriction (Fig. 5A). In the absence of either *ATG16L1* (as expected, (13, 15)) or *WIPI2*, IFNγ-induced growth restriction was reversed and fewer PVs containing 1 parasite were observed (Fig. 5A) while the majority containing ≥8 parasites (Fig. 5A). Thus, as for *ATG16L1*, the role of *WIPI2* in IFNγ-induced inhibition of intracellular replication extends from a cytoplasmic RNA virus to an intravacuolar apicomplexan parasite.

**Figure 5:**
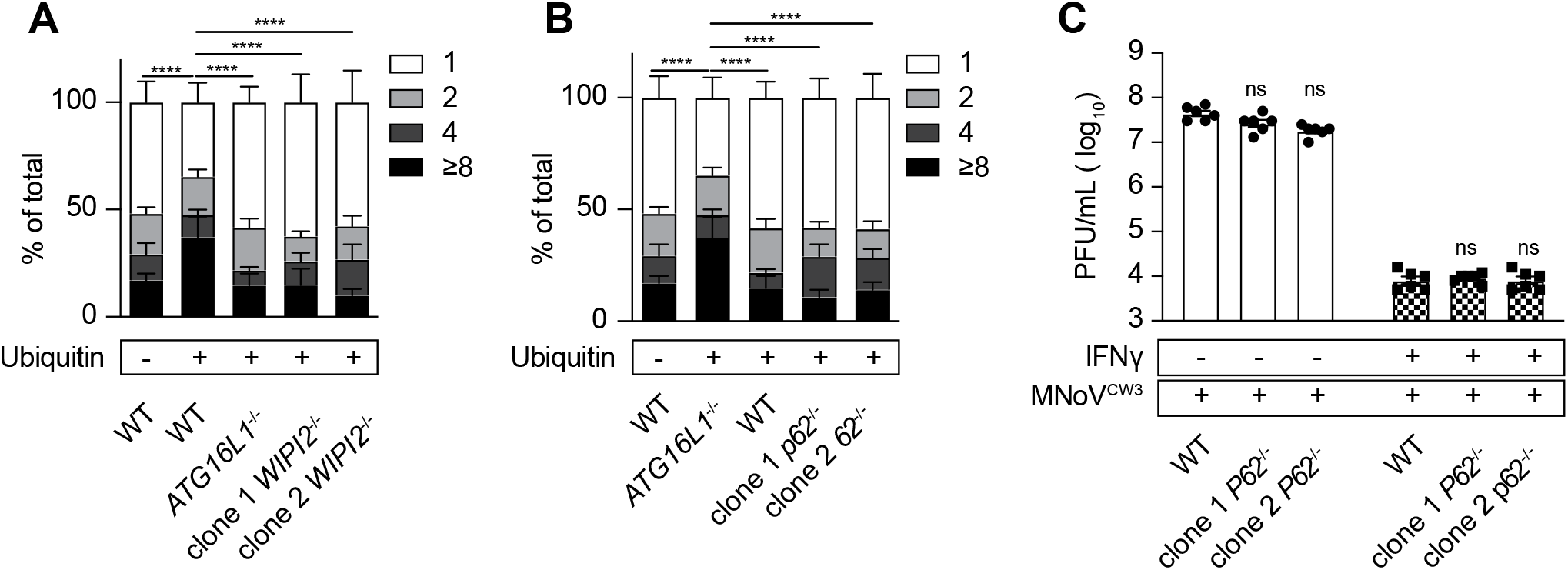
WIPI2 and *P62* are required for IFNγ-induced growth restriction of T. gondii in HeLa cells, *p62* is not required for IFNγ-induced growth restriction of norovirus in BV-2 cells. (A) WT, *ATG16L1*^-/-^and *WIPI2*^-/-^cells were treated with IFNγ and infected with *T. gondii*. Parasites within ubiquitin positive (+) and ubiquitin negative (-) vacuoles were counted by immunofluorescence microscopy. (B) as in (A) with WT, *ATG16L*1^-/-^and *P62*^-/-^cells. (C) Plaque assay of WT and *p62*^-/-^BV-2 cells as in (Fig S1). Values in (A) (B) (C) represent means ± SEM from two to three independent experiments. Significance shown for vacuoles containing 1 parasite per vacuole. *P* value ≤ 0.05 (*), ≤ 0.01(**), ≤0.001 (***), ≤ 0.0001 (****) were considered statistically significant. ns, not significant. ns, not significant. For *T. gondii* assay, *P* value determined by 2-way ANOVA with Tukey’s multiple comparison test; for plaque assay *P* value determined by 2-way ANOVA with Dunnet’s multiple com-parison test.

### *P62* in required for IFNγ-induced inhibition of *T. gondii* in human HeLa cells but not required for IFNγ-induced inhibition of norovirus replication in BV-2 cells

In murine cells, the ubiquitin binding protein P62 is recruited to vacuoles containing *T. gondii* in an IFNγ-dependent manner but is ultimately dispensable for inhibiting replication of the parasite (54). In human cells, P62 is also recruited to parasitophorous vacuoles in an IFNγ-dependent manner (15) but its requirement for the restriction of *T. gondii* replication is unknown. Therefore, we generated clonal *P62*^-/-^HeLa cells (Table S1) and observed that *P62* was required for IFNγ-induced inhibition of *T. gondii* (Fig. 5B) but was not required for IFNγ-induced inhibition of MNoV^CW3^ replication in clonal p62^-/-^BV-2 cells (Fig. 5C)(Table S1). Taken together, these data suggest the requirement for *P62* in IFNγ-mediated control of intracellular pathogens can be species or cell type specific.

## Discussion

Intracellular pathogens present diverse challenges to the immune system because they hijack different aspects of cell biology to their own advantage. IFNγ is a central mediator of innate and adaptive immunity, in part via orchestrating the activities of populations of immune cells. However, IFNγ is also responsible for cell-intrinsic immunity to intracellular pathogens via induction of transcriptional and other cellular pathways that create an environment inside cells that is hostile to invading pathogens. Understanding the molecular and cellular mechanisms for this process is important for defining mechanisms of immunity to infection. We report herein additional cellular genes that regulate intracellular immunity and identify components of the autophagy machinery including *Atg9a* and *Wipi2b* that play an autophagy-independent role in IFNγ-induced intracellular immunity to *T. gondii* and norovirus.

We and others have shown that the mechanisms of IFNγ-induced intracellular immunity to norovirus and *T. gondii* do not require autophagy as a degradative function, but nevertheless require essential *ATG* genes involved in the ubiquitin-like systems that conjugate ATG8 family proteins to PE during autophagy (10–13, 15, 16, 18, 55). The involved proteins include ATG7, which triggers the lipidation of ATG8 family proteins, ATG3 which complexes with ATG8 family proteins to foster their lipidation, and the E3-ligase-like ATG5-12-16L1 complex (10–13, 15, 16, 18, 55), which directs lipidation of ATG8 family proteins to appropriate sub-cellular locations. The ATG8 family comprises LC3 and GABARAP proteins involved in various aspects of autophagy and non-autophagic cellular processes (1, 4, 56, 57). In many of these functions, ATG8 proteins are localized on membranes by lipidation dependent on the ATG5-12-16L1 complex.

It is notable that the cellular membranes targeted by these *ATG* genes are quite distinct for *T. gondii* and norovirus. *T. gondii* survives and replicates via binary fission inside a single membrane-bound vacuole created when it invades the cell. Replication occurs sequestered away from the cytoplasm. In contrast, norovirus replication occurs on the cytoplasmic face of intracellular membranes rearranged into a topologically complex replication complex responsible for orienting viral proteins and nucleic acids for efficient replication and assembly. Nevertheless, these two topologically distinct processes are interdicted by IFNγ in a manner dependent on a common set of *ATG* genes. This *ATG* gene-dependent form of intracellular immunity is triggered by IFNγ binding to its receptor to induce *Stat1* protein phosphorylation, *Irf1* activation and the transcriptional induction of effector proteins (11, 16, 25, 58).

*ATG* genes required for this form of IFNγ-induced immunity provide clues as to the mechanisms involved, but so do the *ATG* genes that are not required. We confirmed the previously reported lack of a role for *Atg14* in IFNγ-induced inhibition of norovirus replication (16). This is a key observation because ATG14 is an essential component of the C1 form of the hetero-tetrameric Class III PI3Kinase complex required for autophagy initiation (1, 3). ATG14 is also not required for control of *T. gondii* in IFNγ-treated human HeLa cells (15). The C1 complex comprises ATG14, VPS15, VPS34 and Beclin 1, while the C2 complex substitutes UVRAG for ATG14 (1, 4, 5). These lipid kinases generate PtdIns(3)P on membranes including the isolation membrane to initiate autophagy. While ATG14 is a component of only one form of the Class III PI3Kinases involved in autophagy, Beclin 1 is a core member of all known autophagy-related Class III PI3Kinases. We report here that the gene *Becn1* was not required for IFNγ-mediated inhibition of norovirus replication. Of note, the specificity of our observations demonstrating a lack of a role for *Atg14* and *Becn1* in IFNγ-induced immunity is supported by the observation that the same cell lines used here were used to show that *Becn1* and *Atg14* are required for protection against IFNγ-induced cell death in BV-2 cells (27). It is notable that there is overlap between the genes required to prevent cell death induced by IFNγ, and those required for IFNγ to inhibit norovirus replication. However, the mechanisms of IFNγ-induced inhibition of norovirus replication and prevention of cell death likely differ because prevention of IFNγ-induced cell death requires *Atg14* and *Becn1* (27), while prevention of norovirus replication did not. Together these data show that the classical PI3K complexes involved in autophagy are not required for this form of intracellular immunity.

### Cellular mechanisms of *ATG* gene-dependent immunity to norovirus and *T. gondii*

The mechanism by which *ATG* proteins inhibit replication of both norovirus and *T. gondii* in IFNγ-treated cells involves the recruitment of IFN-inducible GTPases to the intracellular membranes responsible for supporting pathogen replication (14, 16, 55, 58–60). While the genes encoding sub-families of the interferon-γ-inducible immunity-related GTPase (IRGs) differ between humans and mice (55, 61), IRGs and guanylate-binding proteins (GBPs) in mice, and GBPs in humans (62) are required for IFNγ-induced inhibition of the replication of both norovirus and *T. gondii*, as well as *Chlamydia* (58, 63, 64). In contrast to species variations in IFN-induced GTPases that act downstream of *ATG* proteins, the role of the ATG5-12-16L1 complex is conserved between humans and mice for this form of IFNγ-induced intracellular immunity. There appear to be conserved sequential events that result in targeting norovirus and *T. gondii*-related cellular membranes despite the striking differences in topology between the cellular compartments used by these two pathogens. First, the ATG5-12-16L1 complex is recruited to the relevant membranes followed by deposition of lipidated forms of the ATG8 family proteins. This is followed by recruitment of IFN-inducible GTPases to the target membranes (reviewed in (58)). Less well defined are events prior to ATG5-12-16L1 recruitment and those that occur after recruitment of IFNγ-induced immune effectors to pathogen-related membranes (58).

### New components of *ATG* gene-dependent immunity

We validated the BV-2 cell system for studies of IFNγ-induced inhibition of norovirus via demonstration of roles for *Stat1, Irf1* and the ATG5-12-16L1 complex, confirming the expected role of these genes observed in other systems (11, 16, 23, 25). Validation of BV-2 cells as a system to study *ATG* gene-dependent immunity enabled screening of a set of autophagy-related genes using CRISPR-targeted gene disruption. Mock or IFNγ treatment alone revealed genes essential for cell survival, which were not studied further. Analysis of cells treated with IFNγ and infected with norovirus revealed genes essential for efficient norovirus replication (21, 45–47) and the genes that regulate responses to IFNγ such as *Ufc1* and *Uba5*. We further showed that *Wipi2* and *Atg9a* are required for efficient IFNγ-induced inhibition of norovirus replication; we were able to confirm a role for *WIPI2B* in control of *T. gondii* in addition to norovirus replication. Intriguingly, *Atg9a* was previously found to be dispensable for the localization of *p62*, GBPs, *Irga6* and ubiquitin to the parasitophorous vacuole of *T. gondii* in mouse embryonic fibroblast (17) ; *p62* was also found to be dispensable for inhibiting replication of the parasite in the murine cells (54). However, we found that *P62* was required for IFNγ-induced inhibition of *T. gondii* in HeLa cells, but not for norovirus replication in murine cells. Thus, while there are shared *ATG* genes involved in IFNγ-induced immunity to these two pathogens, there are proteins such as ATG9A and P62, which may play pathogen-or cell type-specific role in these topologically distinct cellular compartments in different experimental systems.

### Role of *Atg9a* in IFNγ-induced *ATG* gene-dependent immunity

*Atg9a* plays its role in autophagy via provision of lipids to developing autophagosomes via recruitment of ATG9A-positive small vesicles (52, 57, 65). While the role for *Atg9a* demonstrated here likely involves these small vesicles, the regulation of *Atg9a* in IFNγ-induced intracellular immunity appears to differ from its regulation during autophagosome formation. In murine cells neither the unc-5-like kinases ULK1 nor ULK2, which are required for initiation of autophagy, are required for IFNγ-mediated inhibition of norovirus replication (16, 57). This is particularly interesting since the role of ATG9A trafficking in autophagy is tightly regulated by ULK1 kinase (57). This suggests that the regulation of *Atg9a* function in IFNγ-induced *ATG* gene-dependent immunity is likely via a regulatory cascade distinct from that utilized to regulate the role of *Atg9a* in autophagy.

Our data that *Atg9a* plays a key role in IFNγ-mediated inhibition of norovirus infection may be explained by the finding that deletion of ATG9 mRNA enhances induction of iNOS protein expression by IFNγ (26). Thus, the role of *Atg9a* might be either direct through action of this protein on pathogen-related intracellular membranes or indirect through changes in IFNγ signaling, or both. Our findings of a negative regulatory role for the UFMylation activity of the UFC1 and UBA5 enzymes is likely due to such an indirect effect on IFNγ signaling, but in this case likely due to the enhancement of IFNγ response via *Ern1*-dependent ER stress responses that occur when UFMylation is inhibited (26).

### Role of WIPI2B in IFNγ-induced *ATG* gene-dependent immunity

The role of *Wipi2b* in our system may inform how the ATG5-12-16L1 complex is recruited to intracellular membranes during IFNγ-induced *ATG* gene-dependent intracellular immunity. WIPI2B binds to both PtdIns(3)P in cellular membranes and to ATG16L1, and is important for intracellular targeting and clearance of *Salmonella enterica serovar* Typhimurium via the recruitment of the Atg12-5-16L1 complex to cellular membranes followed by LC3 lipidation and the formation of autophagosomal membranes to engulf the bacteria (52). We found that amino acids required for WIPI2B to interact with both ATG16L1 and PtdIns(3)P were required for IFNγ-induced inhibition of norovirus replication. One explanation for these findings is that WIPI2B is upstream of the effects of the ATG5-12-16L1 complex, perhaps by binding cellular lipids on target membranes and then recruitment of the ATG5-12-16L1 complex via binding to ATG16L1 as observed for its role in degradative autophagy (52). This finding presents an interesting conundrum in that we found that Beclin 1, an essential component of the Class III Pi3Kinases that generate PtdIns(3)P required for autophagy, is not required for IFNγ-induced *ATG* gene-dependent inhibition of norovirus replication while, WIP2B and the ATG5-12-16L1 complex are required. Interestingly, activation of Beclin 1-depedent processes is not sufficient to target the parasitophorous vacuole inhabited by *T. gondii* for *ATG* gene-dependent immunity (15). These observations open new questions on how pathogen-related membranes become labelled with specific phospholipids and in turn recruit ATG proteins.

## Summary

Data presented here, and significant prior work, demonstrate the existence and physiological importance of a unique IFNγ-induced *ATG* gene-dependent process responsible for creation of a cellular environment that is hostile to the replication of phylogenetically distinct pathogens. This mechanism appears programmable with the participation of a core machinery including WIPI2B, ATG9A, ATG7, ATG5-12-16L1, ATG3 and ATG8 family members rendered pathogen-specific by proteins such as P62 and IFNγ-induced GTPases (11, 16, 58). These new data further support the conclusion that cassettes of *ATG* genes have been leveraged by the immune system to perform cytokine-induced tasks that block replication and clear pathogens as previously proposed (3, 10, 11). This system is programmable via the involvement of a variety of other gene products and focuses on altered intracellular membranes created as diverse pathogens hijack and evade normal cellular functions.

## Materials and Methods

### Cells

BV-2, HEK293T and HeLa cells were cultured in Dulbecco’s Modified Eagle Medium (DMEM, Gibco) with 10% fetal bovine serum (FBS) and 1% HEPES. BV-2 Cas9 cell lines were generated using standard protocols (21). *Stat1*^-/-^, *Irf1*^-/-^, *Atg12*^*-/-*^, *Atg16l1*^*-/-*^, *Wipi2*^-/-^, *p62*^-/-^, *WIPI2*^-/-^and *P62*^-/-^cells were generated by introducing Cas9 and gRNAs into BV2 or HeLa cells by nucleofection. For selection, 5 μg/mL puromycin (Thermofisher) and 5 μg/mL blasticidin (Thermofisher) was added to BV-2 cells.

BV-2 mutant cell lines were generated at the Genome Engineering and iPSC center at Washington University School of Medicine (21). sgRNAs (Table S1) were nucleofected with Cas9 into wild type BV2 cells and clones were screened for indels by sequencing the target region with Illumina MiSeq at approximately 500x coverage. Indel signature frequency was determined using an in-house algorithm (GEiC, Washington University School of Medicine, St. Louis, MO).

For cDNA expression of WIPI2B proteins cells were transduced with lentivirus carrying the gene of interest with an N-terminal 3X Ty1 tag (Table S7) on the pCDH-CMV-MCS-T2A-Puro backbone (CD522A-1, System Biosciences). cDNA expression of ATG proteins has been previously described (21, 26, 27).

### Viruses and viral assays

MNoV^CW3^ (Gen bank accession no. EF014462.1) was generated by transfecting a molecular clone (66) into HEK293T cells (P0 stock), which was passaged on BV-2 cells. After two passages, infected cells were frozen at -80C and thawed, cleared of cellular debris and virus was concentrated by tangential flow filtration. For infection, WT or knockout BV-2 cells were seeded at 10^4^ cells/well of a 96-well plate. After 8 hours, cells were treated with 1U/mL of IFNγ (BioLegend). 16 hours later MNoV^CW3^ was added at an MOI 5.0. Infected cells were harvested at 24 hpi and frozen at -80°C prior to plaque assay. For cell viability CellTiter-Glo reagent (Promega) was added to wells of a 96-well plate, incubated for 10 minutes at room temperature and then cellular ATP content was measured. Viral titers were determined in triplicate by plaque assay on BV-2 cells. BV-2 cells were seeded at 2 x 10^6^ cells/well of a six-well plate and 24 hours later 100uL of 10-fold serially diluted samples were applied to each well for 1 hour with orbital rocking at room temperature. Viral inoculum was aspirated, and 2 ml of warmed MEM containing 10% FBS, 2mM L-Gluatmine, 10 mM HEPES, and 1% methylcellulose was added. Plates were incubated for 48-60 hours prior to visualization after staining with 0.2% crystal violet in 20% ethanol.

### Antibodies and western blots

Cell lysates were run under reducing conditions on an Any kD™ Mini-PROTEAN® TGX Stain-Free™ Protein Gel (BioRad) and imaged using a ChemiDoc imaging system (Biorad). WIPI2B proteins were tagged with three copies of Ty1 on the N terminus (Brookman et al., 1995). Anti-Ty1 antibody (ThermoFisher #MA5-23513) was used at 1:1000. Anti-Mouse HRP (Jackson Immuno Research Laboratories #315-035-003) was used at 1:10,000.

### Autophagy CRISPR-Cas9 subpool generation and screen

1.4×10^7^ BV-2-Cas9 cells (21, 26, 27) were transduced with a lentivirus stock (transduction efficiency 40%) resulting in 6 x10^6^ cells. 36 hours later puromycin was added and transduced cells selected for five days. Five days later 1×10^7^ cells were harvested and DNA isolated for sequencing (QIAamp DNA Blood Midi Kit, Qiagen). For each experimental condition (mock, IFNγ, and IFNγ + MNoV^CW3^) 1×10^7^ cells were seeded in a 15cm^2^ dish. Eight hours later, cells were treated with media or 1 U/ml IFNγ. After 16 hours cells were either mock infected or infected with MNoV^CW3^ at an MOI of 5.0. Cells were harvested 24 hour later for isolation of DNA for sequencing and assessment of cell viability (Trypan blue exclusion). Genomic DNA was sequenced and analyzed (21, 26, 27). Volcano plots were generated using the hypergeometric distribution method (https://github.com/mhegde/volcano_plots) and screen results were analyzed using STARS (https://portals.broadinstitute.org/gpp/public/software/stars).

### *T. gondii* growth restriction assay

HeLa cells were treated with 100 U/ml IFNγ for 24 hours, infected with tachyzoites, and washed 2 hours later to remove extracellular parasites. Cells were fixed in 4% formaldehyde 24 hours later and *T. gondii* localized using antibody against the RH strain tachyzoites (67) and ubiquitin was localized using mouse antibody clone FK2 (04-263; EMD Millipore Corporation). Parasites were enumerated per parasitophorous vacuole (PV) from 30 PV on three individual coverslips from three independent experiments.

### Quantification and Statistical Analysis

Data were analyzed with Prism 7 software (GraphPad Software, San Diego, CA). Volcano plot and Pearson visualizations were generated using R-studio (Integrated Development for R. RStudio, PBC, Boston, MA URL http://www.rstudio.com/).

## Acknowledgments

The authors would like to thank Rob Orchard, Craig Wilen, Yating Wang, Scott Handley, Tim Schaiff, Anne Paredes and John Doench for discussion; Monica Sentmanat in the Genome Engineering and iPSC Center (GEiC) at Washington University School of Medicine for assistance with CRISPR cell line generation. H.W.V. was supported by the National Institutes of Health (NIH) grants U19 AI109725. H.W.V., M.R.M, D.B. and S.H. is supported by U19 AI142784. M.R.M. was supported by U19 AI109725. A.P. was supported by T32 AI716340. J.B. and L.D.S. were supported by NIH grant AI 118426.

## AUTHOR CONTRIBUTIONS

M.R.M. and H.W.V. designed the project and wrote the manuscript. A.O., A.P., M.E.S., J.B., L.D.S., D.B. and S.H. helped edit and review the manuscript. M.R.M, A.O., A.P., M.E.S., J.B., L.D.S., D.B. and S.H. assisted with project design. M.R.M, J.B., A.O., A.P., and M.E.S. designed and/or assisted with experiments and data analysis. All of the authors approved the manuscript.

## Supplemental Materials

**Figure S1.**
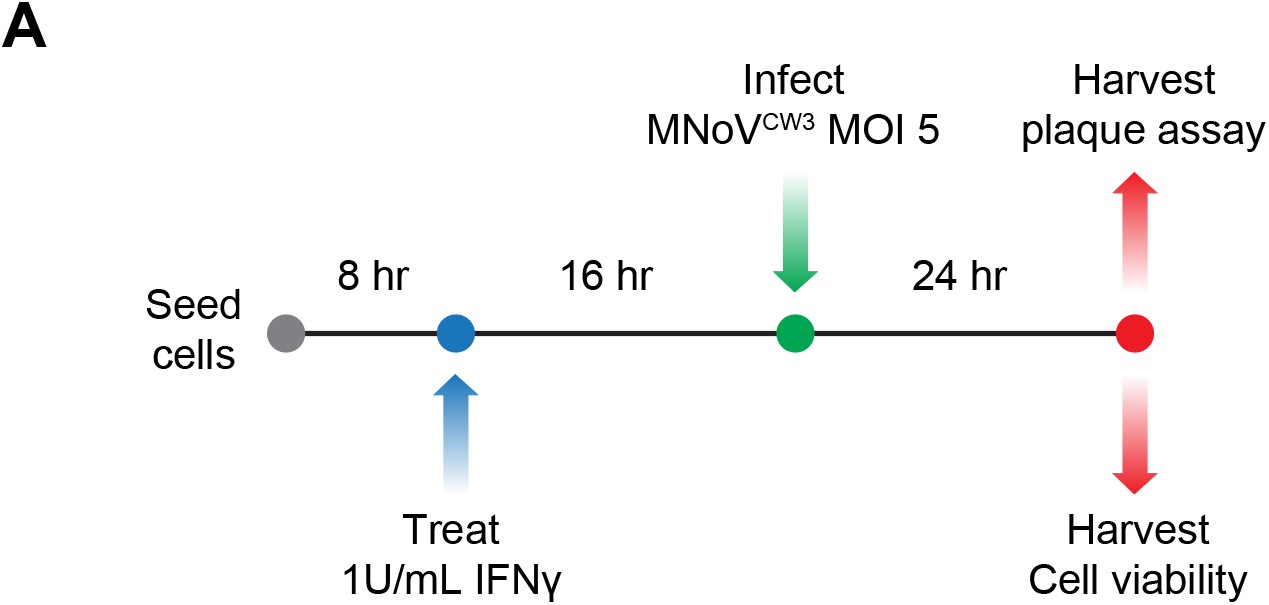
Schematic for seeding, treatment and norovirus infection. (A) Experimental workflow for IFNγ treatment and virus infection in BV-2 cells.

**Figure S2.**
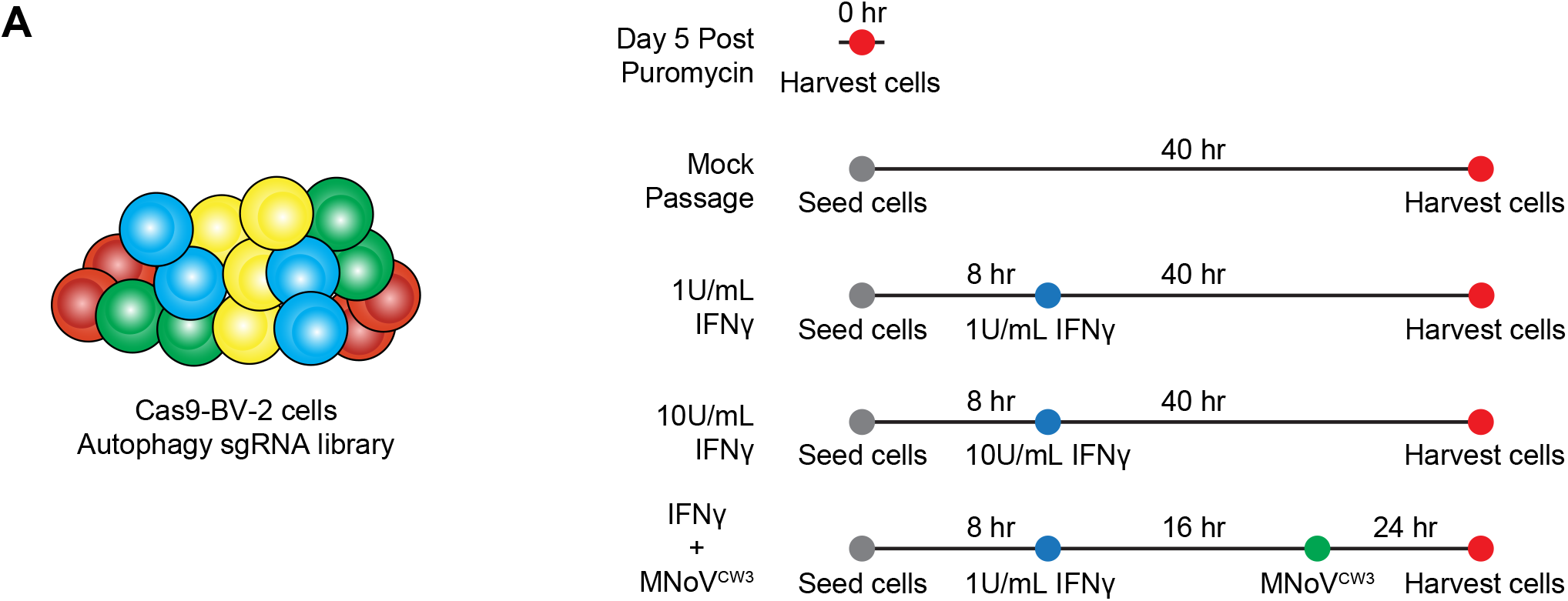
Autophagy CRISPR library screen design. (A) BV2-Cas9 cells transduced with the Autophagy CRISPR library were collected for the following conditions: five days post puromycin selection, mock treatment, 1U/mL IFNγ treatment, 10U/mL IFNγ treatment or 1U/mL IFNγ treatment + MNoV^CW3^.

## Supplemental Datasets

**Supplemental Tables 1. NGS validation data. Related to Figure 1**,**3**,**4**,**5**. List of sgRNA sequences used in this report and validation of editing efficiencies by NGS.

**Supplemental Table 2. Autophagy CRISPR library. Related to Figure 2**. Annotation of sgRNAs and associated mouse genes in the autophagy CRISPR library.

**Supplemental Tables 3. Genes enriched and depleted for passage of BV-2 cells results. Related to Figure 2**. Average LFC and STARS analysis for the CRISPR screen.

**Supplemental Table 4. Genes depleted during passage of BV-2 cells compared to Project Achilles. Related to Figure 2**. Candidate essential genes with FDR < 0.1 compared to Achilles Project annotations.

**Supplemental Tables 5. Genes enriched and depleted in 1U/mL or 10U/mL IFNγ treated cells. Related to Figure 2**. Average LFC and STARS analysis for the CRISPR screen.

**Supplemental Tables 6. Genes enriched and depleted in 1U/mL IFN + MNoV**^**CW3**^ **infected cells. Results related to Figure 3**. Average LFC and STARS analysis for the CRISPR screen.

**Supplemental Table 7. Supplemental sequences. Related to Methods**. cDNA sequences for *Wipi2*^-/-^complementation studies.

